# Evidence for distinct networks underlying symptom clusters of posttraumatic stress disorder

**DOI:** 10.1101/2024.12.31.630851

**Authors:** Katherina K. Hauner, Aileen Chau, Frank Krueger, Barry Gordon, Jordan Grafman

**Affiliations:** Department of Medical Social Sciences, Feinberg School of Medicine, Northwestern University, Chicago, IL; Department of Neurology, Feinberg School of Medicine, Northwestern University, Chicago, IL; Brain Injury Research Program, Shirley Ryan AbilityLab, Chicago, IL; School of Systems Biology, George Mason University, Fairfax, VA; Department of Psychology, George Mason University, Fairfax, VA; Cognitive Neurology/Neuropsychology Division of the Department of Neurology, Johns Hopkins University School of Medicine, Baltimore, MD; Cognitive Science Department, Johns Hopkins University, Baltimore, MD; Department of Physical Medicine and Rehabilitation, Feinberg School of Medicine, Northwestern University, Chicago, IL

**Keywords:** Amygdala, Hippocampus, PTSD, Traumatic Brain Injury, Ventromedial Prefrontal Cortex, Voxel-Based Lesion-Symptom Mapping

## Abstract

**Background:** Clinical and psychometric evidence has long supported a multidimensional model of PTSD, with symptom subcategories derived from factor analytic methods. Although research on the biological bases of PTSD as a unitary construct is profuse, comparatively few studies have examined the neural mechanisms underlying subcategories of PTSD symptoms. The present study aimed to provide the first evidence of causal relationships between brain structure and PTSD symptom subcategories, using a lesion-behavior mapping approach.

**Methods:** Using a group of male combat veterans with focal penetrating traumatic brain injuries (*n* = 177), we determined the effects of focal damage on the PTSD symptom subcategories of hyperarousal, avoidance, and re-experiencing.

**Results:** Our findings revealed two distinct networks that underlie symptom subcategories of PTSD: (1) an amygdala-ventromedial prefrontal cortex network underlying hyperarousal and avoidance symptoms, in which amygdala damage acts as a risk factor for the development of these symptoms, while vmPFC damage acts as a protective factor against the same symptoms; and (2) a hippocampal network underlying re-experiencing and avoidance, in which hippocampal damage acts as a protective factor against these symptoms.

**Conclusions:** The present study provides novel insights regarding the causal role of key brain regions in the heterogeneous expression of PTSD symptoms. Results not only contribute to a more complete picture of the neural mechanisms underlying PTSD, but may also aid in the future development of individualized therapeutic strategies that target specific symptom profiles.

## Introduction

Approximately 12 million Americans are diagnosed with posttraumatic stress disorder (PTSD) [1], a debilitating condition including symptoms of hypervigilance, trauma cue avoidance, and intrusive memories [2]. Although PTSD is normally regarded as a categorical construct, clinical and psychometric evidence has long supported a multidimensional model [3–10] with symptom clusters derived from factor analytic methods. Multidimensional measures of PTSD symptoms have identified a varying number of symptom clusters that typically range from two to five factors and include re-experiencing, hyperarousal, emotional numbing, and avoidance [11]. While symptom clusters have been applied over decades of PTSD research, remarkably little attention has been given to determining their biological bases. In particular, it is unknown whether symptom clusters are subserved by unique neural networks or whether they map onto a distributed network associated with the categorical diagnosis of PTSD. Identifying networks that underlie PTSD symptom clusters may be helpful in guiding the development of future treatments [12], in which individualized approaches could be used to target distinct neurobiological profiles.

Research on the general pathophysiology of PTSD has identified contributions from regions involved in emotional processing and memory, such as amygdala, ventromedial prefrontal cortex (vmPFC), insula, and hippocampus [13–21]. In PTSD patients, amygdala volume is typically decreased [17, 22, 23]; however, this observation provides no information regarding the direction of this relationship. Our prior research addressed the issue of directionality by applying voxel-based lesion symptom mapping in a group of Vietnam veterans with focal brain injuries. These findings revealed that damage to the amygdala and vmPFC is associated with a decrease in PTSD incidence, thus suggesting that this lesions in these regions may serve as a protective factor against the disorder [19]. Functional neuroimaging studies have reliably demonstrated amygdala hyperreactivity during traumatic stimulus presentation, as well as correlations between PTSD severity and hyperreactivity in amygdala and insula (as well as hypoactivity in vmPFC) [e.g., 13-15, 24, 25]. PTSD researchers have also examined the hippocampus, with a large number of studies indicating that PTSD patients have low hippocampal volume, even in comparison to trauma-exposed participants without PTSD (suggesting that this result is not purely due to the experience of the trauma itself) [13, 21, 23, 26–28]. Studies using task-based fMRI in PTSD patients have also revealed abnormal hippocampal activity during trauma-related tasks [29–32]. While the causal direction of these results is again unclear, our own prior findings suggest that hippocampal damage itself may not influence PTSD diagnosis, based on results using lesion-mapping approaches among veterans with focal damage to this region [19].

While these studies have identified neural mechanisms underlying PTSD as a unitary construct, far less is known regarding mechanisms of symptom clusters. Of the few studies on this topic, the majority have focused on the role of the amygdala in hyperarousal. These studies have reported associations between increased hyperarousal in PTSD patients and reduced amygdala volume [33–35], reduced amygdala-mPFC connectivity [35, 36], and increased amygdala activation (in response to threatening stimuli) [35, 37]. In addition to studies on hyperarousal, one recent study reported an association between increased right amygdala volume and avoidance symptoms in a sample of women with PTSD [34] and another identified a correlation between decreased amygdala volume and increased re-experiencing symptoms [38]. A recent review comparing neurocircuitry of PTSD symptom clusters between human and animal studies underscored the suggestion that intrusive thoughts may be caused by “under-modulation” of the limbic system by prefrontal circuits, as supported by multiple studies across species demonstrating amygdala hyperactivity and prefrontal hypoactivity following trauma[35].

Research on prefrontal regions involved in PTSD symptom clusters has largely focused on vmPFC [35]; however, in comparison to studies on the amygdala, research on the vmPFC has produced more varied results, with this region implicated in several PTSD symptom clusters, including hyperarousal, re-experiencing, and avoidance [35, 37, 39–42]. As mentioned above, some researchers subscribe to the notion that activity in this region is negatively correlated with symptoms of intrusive thoughts [35] and hyperarousal [39], while others have reported a positive association between vmPFC activity and hyperarousal/re-experiencing symptoms [37]. Tursich et al. identified a link between increased avoidance and reduced vmPFC connectivity (with anterior cingulate cortex), but only for a subset of symptoms within this cluster [42].

Other studies have reported associations between reduced vmPFC-hippocampal connectivity and increased hyperarousal, re-experiencing, and avoidance symptoms [41], as well as disrupted hippocampal resting-state connectivity and increased re-experiencing symptoms [40]. In a recent report, Clancy and colleagues (2024) examined relationships between hippocampal resting-state connectivity and trauma-related intrusive memories (a re-experiencing symptom) [43], and found associations between regional co-activation patterns of the hippocampal-cortical network and intrusive memory features. In a separate study, Crombie and colleagues (2021) examined a sample of women with PTSD, and found that re-experiencing symptom severity was associated with decreased cortical thickness in the right parahippocampal and medial temporal gyrus as well as left inferior and middle temporal gyrus, and that active avoidance symptom severity was associated with increased cortical thickness in posterior cingulate and occipital regions [34]. Avoidance symptoms have been further connected to the hippocampus during fear-related tasks [35], including in a study that observed positive correlations between avoidance symptom severity and increased hippocampal activity during a fear conditioning paradigm [44]. Finally, a handful of studies on PTSD symptom clusters have examined the insula, with one study identifying an association between hyperarousal and decreased insular connectivity [within the salience network; 42], and other studies reporting links between disrupted connectivity and re-experiencing/avoidance symptoms [36, 41].

While the studies reviewed above point to the amygdala, vmPFC, hippocampus, and insula as key regions implicated in the pathophysiology of PTSD symptom clusters, research on this topic remains sparse and unreplicated. Furthermore, the above studies largely rely on neuroimaging methods, which offer limited information regarding causal relationships between brain structure and function [45]. In order to determine whether a particular region plays a causal role in PTSD symptom development, or whether individual symptom clusters themselves lead to dysregulated brain activity, research relating focal brain damage to PTSD symptomatology is crucial. If links between lesion location and PTSD symptom clusters were to be established, this information would provide valuable insight regarding the causal role of brain regions in the heterogeneous expression of this disorder.

To address this question, we examined a group of veterans from the Vietnam Head Injury Study (VHIS) who sustained focal penetrating traumatic brain injuries (TBI) during combat. We aimed to determine the effects of focal damage in a priori defined regions on PTSD symptom clusters. Specifically, we examined the effects of damage to amygdala, vmPFC, hippocampus, and insula on *hyperarousal*, *avoidance*, and *re-experiencing* symptoms of PTSD. Based on prior results linking hyperarousal to amygdala [33, 36, 37] and vmPFC [37, 39, 41], we hypothesized that damage to these regions would be associated with this particular symptom cluster. Furthermore, given evidence supporting a specific role for the hippocampus in intrusive traumatic memories and PTSD-related flashbacks [46, 47], as well as reported associations between hippocampal connectivity and trauma re-experiencing [36], we predicted that volume loss in this region would be related to the re-experiencing symptom cluster. Given the noted inconsistency in previous findings, our hypotheses were non-directional.

## Methods and Materials

### Participants

Participants included male combat veterans drawn from the Vietnam Head Injury Study (VHIS), a multi-phase longitudinal study conducted at the National Institute of Neurological Disorders and Stroke (NINDS). The VHIS has provided the rare opportunity to explore brain-behavior relationships in a large cohort of veterans with focal penetrating TBI as well as a matched control group of veterans with no history of brain injury. For a detailed description of the VHIS, see Raymont et al., 2011 [48].

The present study involved participants who completed Phase III (2002–2005) of the VHIS (*N* = 254). Of these participants, 231 completed all CAPS testing and CT scanning procedures (described below). Our results are thus based on data collected from these 231 participants (Table 1), and include both TBI patients (*n* = 177) and matched control participants (*n* = 54). The Institutional Review Board at the NINDS in Bethesda (MD) approved study procedures, and all participants provided written consent.

**Table 1.**
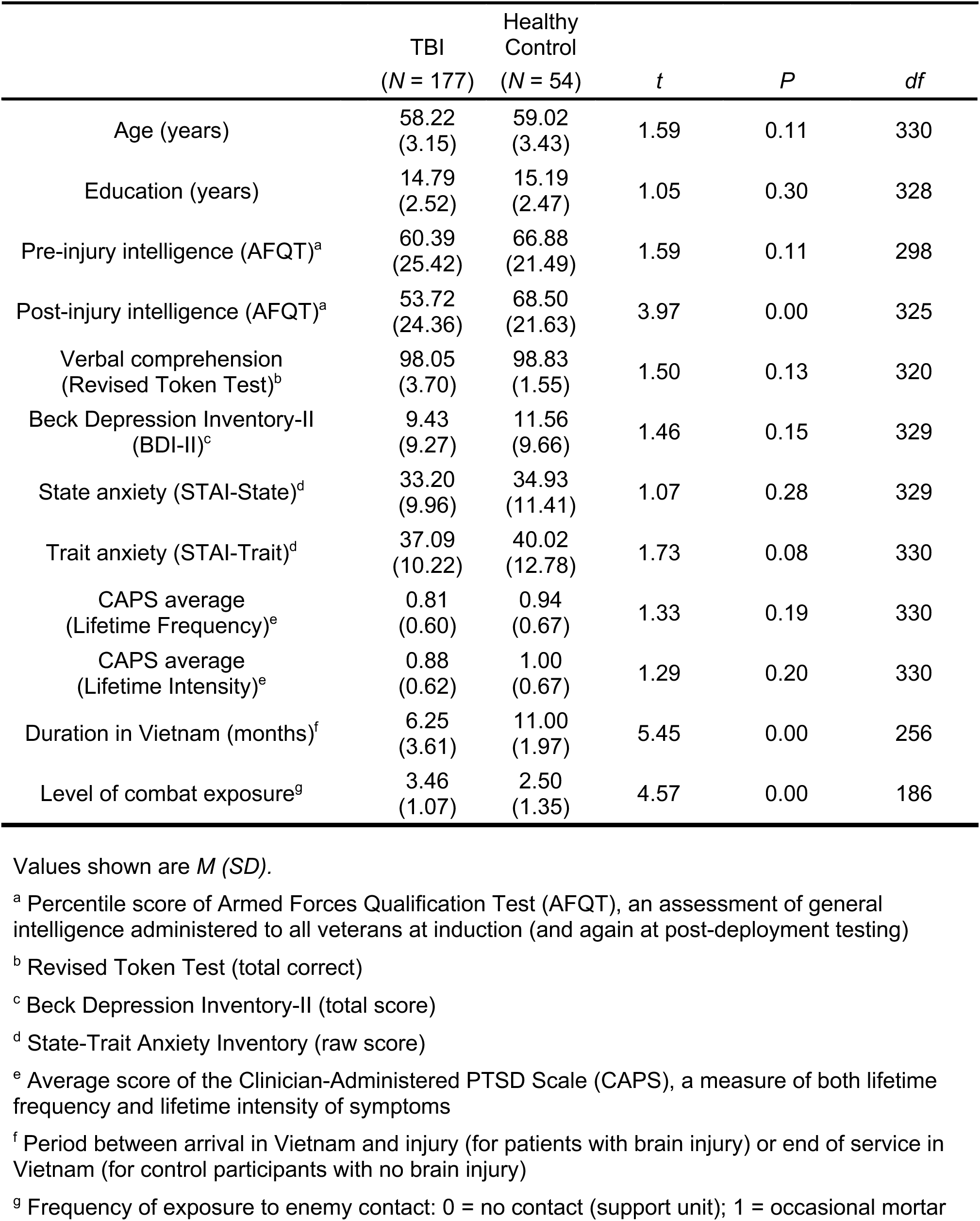
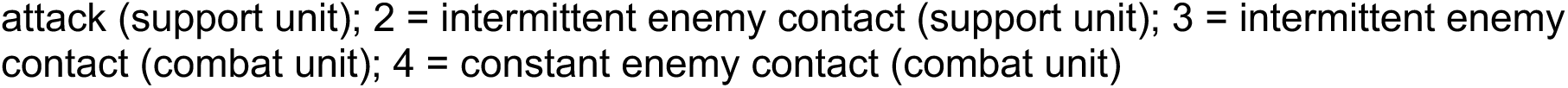
Demographic and neuropsychological measures for all patients with traumatic brain injury (TBI) and healthy participants with no brain injury (healthy control).

|  | TBI<br>(*N* = 177) | Healthy<br>Control<br>(*N* = 54) | *t* | *P* | *df* |
| --- | --- | --- | --- | --- | --- |
| Age (years) | 58.22<br>(3.15) | 59.02<br>(3.43) | 1.59 | 0.11 | 330 |
| Education (years) | 14.79<br>(2.52) | 15.19<br>(2.47) | 1.05 | 0.30 | 328 |
| Pre-injury intelligence (AFQT)<sup>a</sup> | 60.39<br>(25.42) | 66.88<br>(21.49) | 1.59 | 0.11 | 298 |
| Post-injury intelligence (AFQT)<sup>a</sup> | 53.72<br>(24.36) | 68.50<br>(21.63) | 3.97 | 0.00 | 325 |
| Verbal comprehension<br>(Revised Token Test)<sup>b</sup> | 98.05<br>(3.70) | 98.83<br>(1.55) | 1.50 | 0.13 | 320 |
| Beck Depression Inventory-II<br>(BDI-II)<sup>c</sup> | 9.43<br>(9.27) | 11.56<br>(9.66) | 1.46 | 0.15 | 329 |
| State anxiety (STAI-State)<sup>d</sup> | 33.20<br>(9.96) | 34.93<br>(11.41) | 1.07 | 0.28 | 329 |
| Trait anxiety (STAI-Trait)<sup>d</sup> | 37.09<br>(10.22) | 40.02<br>(12.78) | 1.73 | 0.08 | 330 |
| CAPS average<br>(Lifetime Frequency)<sup>e</sup> | 0.81<br>(0.60) | 0.94<br>(0.67) | 1.33 | 0.19 | 330 |
| CAPS average<br>(Lifetime Intensity)<sup>e</sup> | 0.88<br>(0.62) | 1.00<br>(0.67) | 1.29 | 0.20 | 330 |
| Duration in Vietnam (months)<sup>f</sup> | 6.25<br>(3.61) | 11.00<br>(1.97) | 5.45 | 0.00 | 256 |
| Level of combat exposure<sup>g</sup> | 3.46<br>(1.07) | 2.50<br>(1.35) | 4.57 | 0.00 | 186 |

Values shown are *M* (*SD*).
<sup>a</sup> Percentile score of Armed Forces Qualification Test (AFQT), an assessment of general intelligence administered to all veterans at induction (and again at post-deployment testing)
<sup>b</sup> Revised Token Test (total correct)
<sup>c</sup> Beck Depression Inventory-II (total score)
<sup>d</sup> State-Trait Anxiety Inventory (raw score)
<sup>e</sup> Average score of the Clinician-Administered PTSD Scale (CAPS), a measure of both lifetime frequency and lifetime intensity of symptoms
<sup>f</sup> Period between arrival in Vietnam and injury (for patients with brain injury) or end of service in Vietnam (for control participants with no brain injury)
<sup>g</sup> Frequency of exposure to enemy contact: 0 = no contact (support unit); 1 = occasional mortar
attack (support unit); 2 = intermittent enemy contact (support unit); 3 = intermittent enemy contact (combat unit); 4 = constant enemy contact (combat unit)

### Neuropsychological Tests

All participants in Phase III of the VHIS completed an extensive battery of neuropsychological tests over 5–7 days. As shown in tables 1 and 2, we report the following subset: the Armed Forces Qualification Test [AFQT; 49], used to compare pre-and post-injury general intelligence; the Revised Token Test [50], which measures verbal comprehension; the Beck Depression Inventory [BDI-II; 51], to assess symptoms of depression; and the State-Trait Anxiety Inventory [STAI; 52], to evaluate “state anxiety” (momentary) and “trait anxiety” (chronic).

**Table 2.**
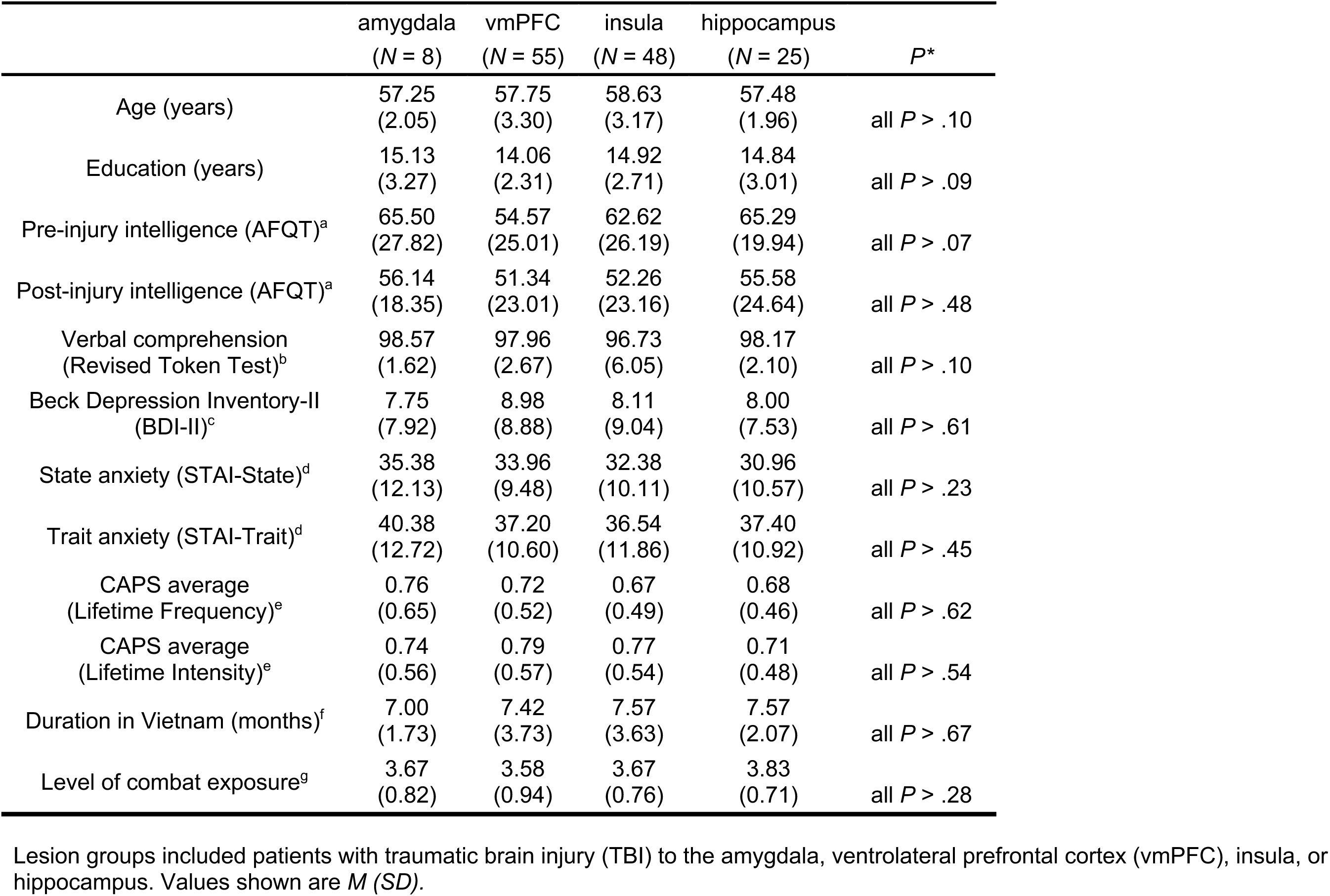
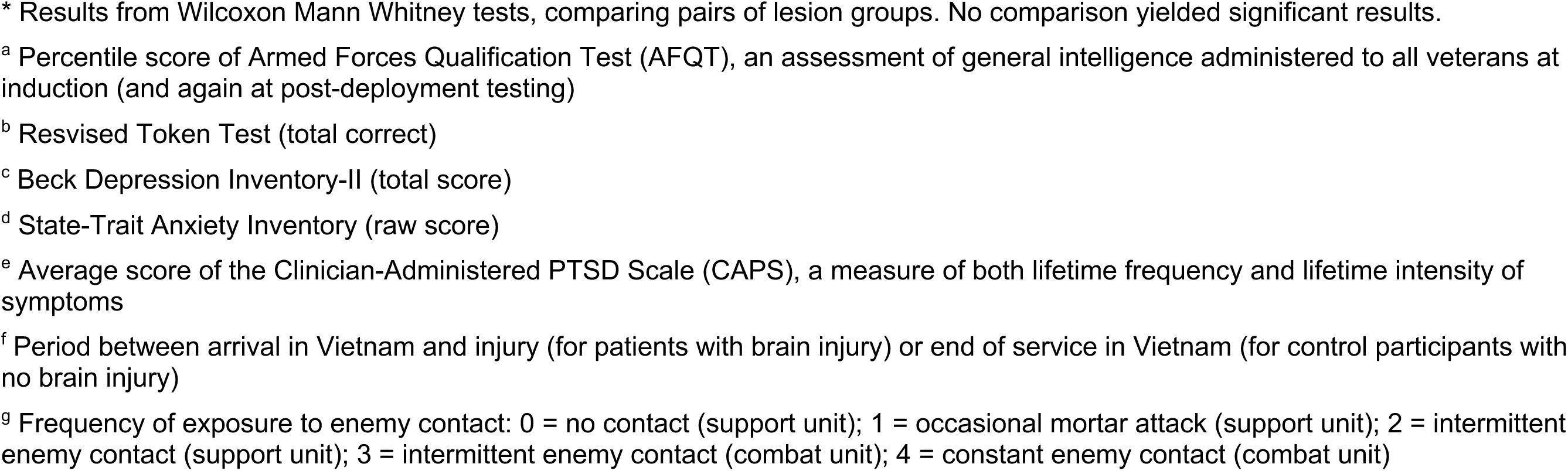
Demographic and neuropsychological measures for patients in lesion groups.

|  | amygdala<br>(*N* = 8) | vmPFC<br>(*N* = 55) | insula<br>(*N* = 48) | hippocampus<br>(*N* = 25) | *P* * |
| --- | --- | --- | --- | --- | --- |
| Age (years) | 57.25<br>(2.05) | 57.75<br>(3.30) | 58.63<br>(3.17) | 57.48<br>(1.96) | all *P* > .10 |
| Education (years) | 15.13<br>(3.27) | 14.06<br>(2.31) | 14.92<br>(2.71) | 14.84<br>(3.01) | all *P* > .09 |
| Pre-injury intelligence (AFQT)<sup>a</sup> | 65.50<br>(27.82) | 54.57<br>(25.01) | 62.62<br>(26.19) | 65.29<br>(19.94) | all *P* > .07 |
| Post-injury intelligence (AFQT)<sup>a</sup> | 56.14<br>(18.35) | 51.34<br>(23.01) | 52.26<br>(23.16) | 55.58<br>(24.64) | all *P* > .48 |
| Verbal comprehension<br>(Revised Token Test)<sup>b</sup> | 98.57<br>(1.62) | 97.96<br>(2.67) | 96.73<br>(6.05) | 98.17<br>(2.10) | all *P* > .10 |
| Beck Depression Inventory-II<br>(BDI-II)<sup>c</sup> | 7.75<br>(7.92) | 8.98<br>(8.88) | 8.11<br>(9.04) | 8.00<br>(7.53) | all *P* > .61 |
| State anxiety (STAI-State)<sup>d</sup> | 35.38<br>(12.13) | 33.96<br>(9.48) | 32.38<br>(10.11) | 30.96<br>(10.57) | all *P* > .23 |
| Trait anxiety (STAI-Trait)<sup>d</sup> | 40.38<br>(12.72) | 37.20<br>(10.60) | 36.54<br>(11.86) | 37.40<br>(10.92) | all *P* > .45 |
| CAPS average<br>(Lifetime Frequency)<sup>e</sup> | 0.76<br>(0.65) | 0.72<br>(0.52) | 0.67<br>(0.49) | 0.68<br>(0.46) | all *P* > .62 |
| CAPS average<br>(Lifetime Intensity)<sup>e</sup> | 0.74<br>(0.56) | 0.79<br>(0.57) | 0.77<br>(0.54) | 0.71<br>(0.48) | all *P* > .54 |
| Duration in Vietnam (months)<sup>f</sup> | 7.00<br>(1.73) | 7.42<br>(3.73) | 7.57<br>(3.63) | 7.57<br>(2.07) | all *P* > .67 |
| Level of combat exposure<sup>g</sup> | 3.67<br>(0.82) | 3.58<br>(0.94) | 3.67<br>(0.76) | 3.83<br>(0.71) | all *P* > .28 |

Lesion groups included patients with traumatic brain injury (TBI) to the amygdala, ventrolateral prefrontal cortex (vmPFC), insula, or hippocampus. Values shown are *M* (*SD*).
\* Results from Wilcoxon Mann Whitney tests, comparing pairs of lesion groups. No comparison yielded significant results.
<sup>a</sup> Percentile score of Armed Forces Qualification Test (AFQT), an assessment of general intelligence administered to all veterans at induction (and again at post-deployment testing)
<sup>b</sup> Revised Token Test (total correct)
<sup>c</sup> Beck Depression Inventory-II (total score)
<sup>d</sup> State-Trait Anxiety Inventory (raw score)
<sup>e</sup> Average score of the Clinician-Administered PTSD Scale (CAPS), a measure of both lifetime frequency and lifetime intensity of symptoms
<sup>f</sup> Period between arrival in Vietnam and injury (for patients with brain injury) or end of service in Vietnam (for control participants with no brain injury)
<sup>g</sup> Frequency of exposure to enemy contact: 0 = no contact (support unit); 1 = occasional mortar attack (support unit); 2 = intermittent enemy contact (support unit); 3 = intermittent enemy contact (combat unit); 4 = constant enemy contact (combat unit)

In addition, participants completed the Clinician-Administered PTSD Scale-Diagnostic Version IV [CAPS IV; 11]. As the standard assessment tool for detailed measurement of PTSD symptoms, the CAPS assesses both frequency and intensity of PTSD symptoms. For the present study, data were collected on lifetime frequency and lifetime intensity, with each symptom rated on a 0–4 scale (where 0 represented “none” and 4 represented “most or all of the time” or “extreme”). The CAPS-IV divides all 17 PTSD symptoms into the following three clusters: (a) *re-experiencing* (5 symptoms), (b) *avoidance/numbing* (7 symptoms), and (c) *hyperarousal* (5 symptoms). Critically, for all participants, the CAPS was administered with combat-related exposure as the traumatic event (Criterion A). Participants who responded “0” to every question were excluded (*n* = 11), to account for potential non-compliance, fatigue, or lack of attention/comprehension. Finally, to assess the extent of trauma exposure, we also collected data on duration of military service in Vietnam and level of exposure to enemy contact (as listed in tables 1 and 2).

### CT Acquisition and Lesion Identification

Computer Tomography (CT) scans were acquired during Phase III of the VHIS (2002– 2005) at Bethesda Naval Hospital, using a General Electric Medical System Light Speed Plus scanner in helical mode. CT (rather than MRI) was used because a large number of participants had retained intracranial metal shrapnel and surgical clips due to their injuries. Structural neuroimaging data were reconstructed with an in-plane voxel size of 0.4 x 3 x 0.4 mm^3^, overlapping slice thickness of 2.5 mm, and 1-mm slice interval. Lesion location and volume loss were obtained using the interactive analysis of brain lesions (ABLe) software implemented in MEDx v3.44 (Medical Numerics) [53, 54]. Each lesion was manually outlined in every slice in native space by a trained neuropsychiatrist. Lesion areas were then reviewed by the senior author (JG), who was blind to neuropsychological evaluation results. Lesion volume was calculated by summing all traced areas and multiplying by slice thickness. Scans were spatially normalized to a CT template brain image in MNI space [55]. Using a 12-parameter affine fit, spatial normalization was performed with the automated image registration (AIR) algorithm [56] to improve registration accuracy.

### Lesion Group Selection

Participants were selected for inclusion in four lesion groups: amygdala, vmPFC, hippocampus, and insula (see Figure 1). Because the amygdala, hippocampus, and insula are pre-defined AAL structures, it was unnecessary to specify criteria for these regions. We defined the vmPFC ROI according to AAL structures within a range of MNI coordinates [19, 57–59]. The vmPFC ROI included portions of the following AAL structures, bounded by the MNI coordinates z ≤ 1, −20 ≤ x < 0 (left hemisphere), 0 ≤ x ≤ 20 (right hemisphere): superior frontal gyrus (orbital), superior frontal gyrus (medial), middle frontal gyrus (orbital), inferior frontal gyrus (orbital), olfactory cortex, gyrus rectus, anterior cingulate and paracingulate gyri. These criteria defined a region comprising the ventral portion of the medial prefrontal cortex (below the level of the genu of the corpus callosum) and the medial portion of the orbital surface (approximately the medial one-third of the orbitofrontal cortex in each hemisphere), as well as subjacent white matter.

**Figure 1.**
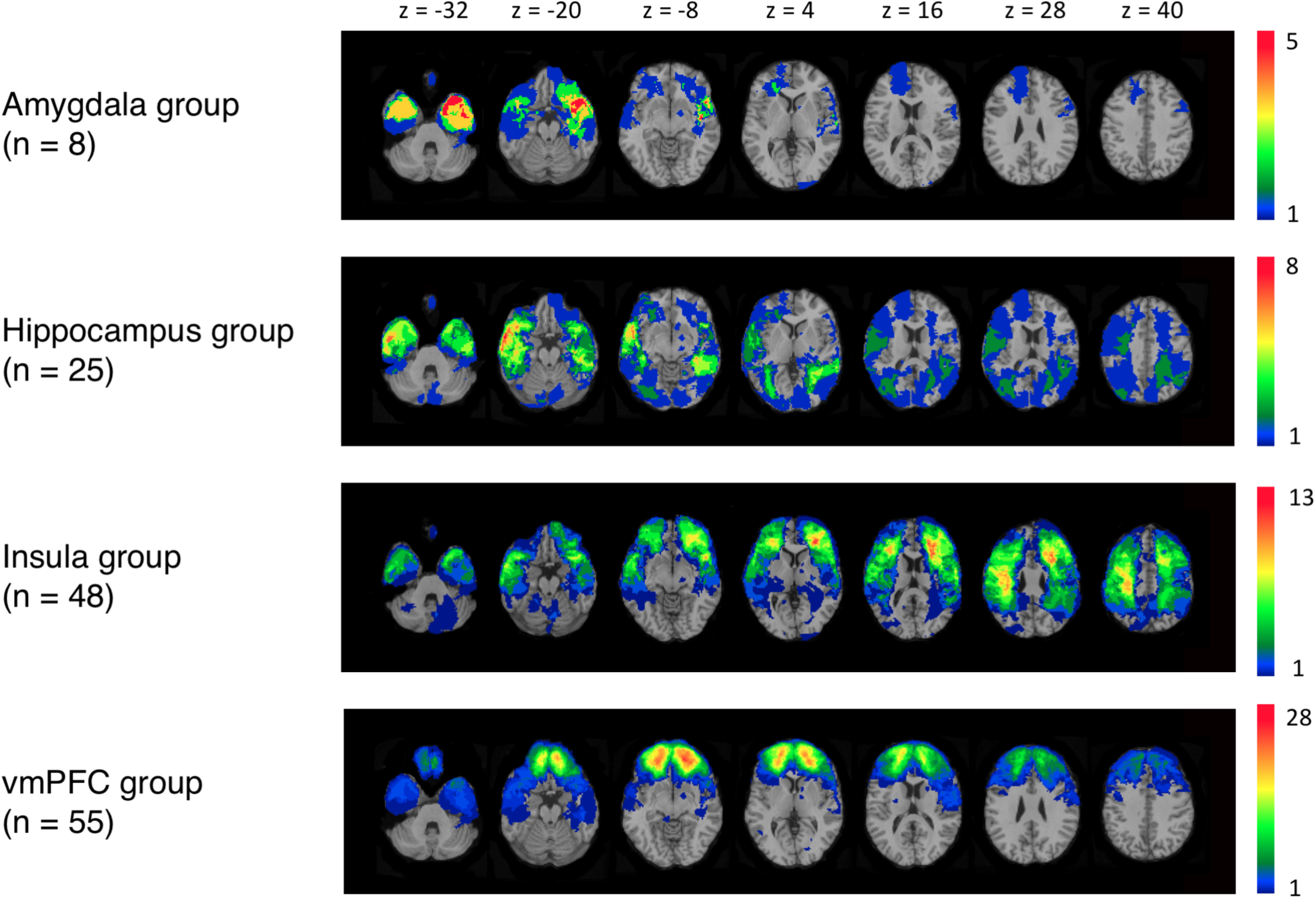
Lesion overlay map of TBI patients (*N* = 177). Values shown in white indicate the *z* coordinates (MNI) of each axial slice. Warmer colors indicate greater lesion overlap (units: number of patients with lesion[s] in this region). All images are depicted in radiological convention (i.e., right hemisphere is on the reader’s left).

Lesion size was defined as the percentage of structure that was damaged within each ROI. For our primary set of analyses, in which we examined correlations between lesion size of each ROI and PTSD symptom cluster, patients were included in a lesion group if both of the following criteria were met: (1) > 0% damage to the ROI for that lesion group, and (2) < 15% damage to any of the three additional ROIs [as in our previous methods; 19, 60, 61]. No minimum ROI lesion size was selected, in order to capture the maximum possible variance for this variable, given the use of correlational tests. For subsequent analyses involving paired comparisons between lesion groups, this criterion was adjusted to ensure that no participant met criteria for more than one lesion group. In these analyses, a threshold of 15% lesion size was used as the inclusionary criterion, while still excluding patients with lesions > 15% in the three additional ROIs.

### Statistical Analysis

Behavioral data analyses centered on the CAPS-IV PTSD symptom clusters as the main outcome variables. For each symptom cluster, lifetime frequency and lifetime intensity were assessed. All outcome variables were examined for non-normality using Shapiro-Wilk tests; significant results indicated that non-parametric testing was necessary. Outcome variables were Winsorized to better approximate a normal distribution curve without removing data. To examine the relationship between lesion volume and symptom clusters, Spearman’s rank correlations (all two-tailed) were performed for each lesion group. Next, in order to directly compare lesion groups, a series of Wilcoxon Mann Whitney paired comparisons was conducted. These analyses tested specific hypotheses that damage to amygdala and vmPFC would more likely be associated with hyperarousal symptoms than damage to other regions, and that damage to the hippocampus, in particular, would be associated with re-experiencing symptoms. All analyses were performed using R version 3.5.0 [62].

## Results

### Effects of Lesion Size

As hypothesized, correlational analyses revealed a positive association between amygdala lesion volume and hyperarousal symptoms, on measures of symptom frequency (*r* = 0.70, *P* = 0.05) and intensity (*r* = 0.72, *P* = 0.04). Amygdala lesion volume was also positively associated with the symptom cluster of avoidance, again for both frequency (*r* = 0.89, *P* = 0.003) and intensity (*r* = 0.89, *P* = 0.003) of symptoms. Lesion volume for vmPFC was also associated with hyperarousal; however, the relationship direction was reversed, with increased lesion volume associated with decreased symptom frequency (*r* = -0.28, *P* = 0.04). Together, these results indicate that lesion size in both regions may be of clinical significance in patients with PTSD, with vmPFC and amygdala playing contrasting roles in their effects on symptom frequency.

### Lesion Group Comparisons

To test hypotheses regarding specific lesion group comparisons (described above), a series of Wilcoxon Mann Whitney tests was performed.

Results indicated that patients with amygdala lesions showed greater frequency of avoidance symptoms than patients with vmPFC lesions (*U* = 40.5, *P* = 0.04), as illustrated in Figure 2. In addition, intensity of avoidance symptoms was similarly higher for patients with amygdala vs. vmPFC lesions (*U* = 40.5, *P* = 0.04). Finally, patients with vmPFC lesions reported reduced hyperarousal intensity in comparison to those with insular lesions (*U* = 102.5, *P* = 0.03). These results indicate that vmPFC lesions were particularly associated with *decreased* hyperarousal and avoidance symptoms. It bears mentioning that participants with vmPFC lesions consistently exhibited the lowest scores on all measures of hyperarousal and avoidance (i.e., avoidance frequency, avoidance intensity, hyperarousal frequency, hyperarousal intensity) in comparison to any other lesion group.

**Figure 2.**
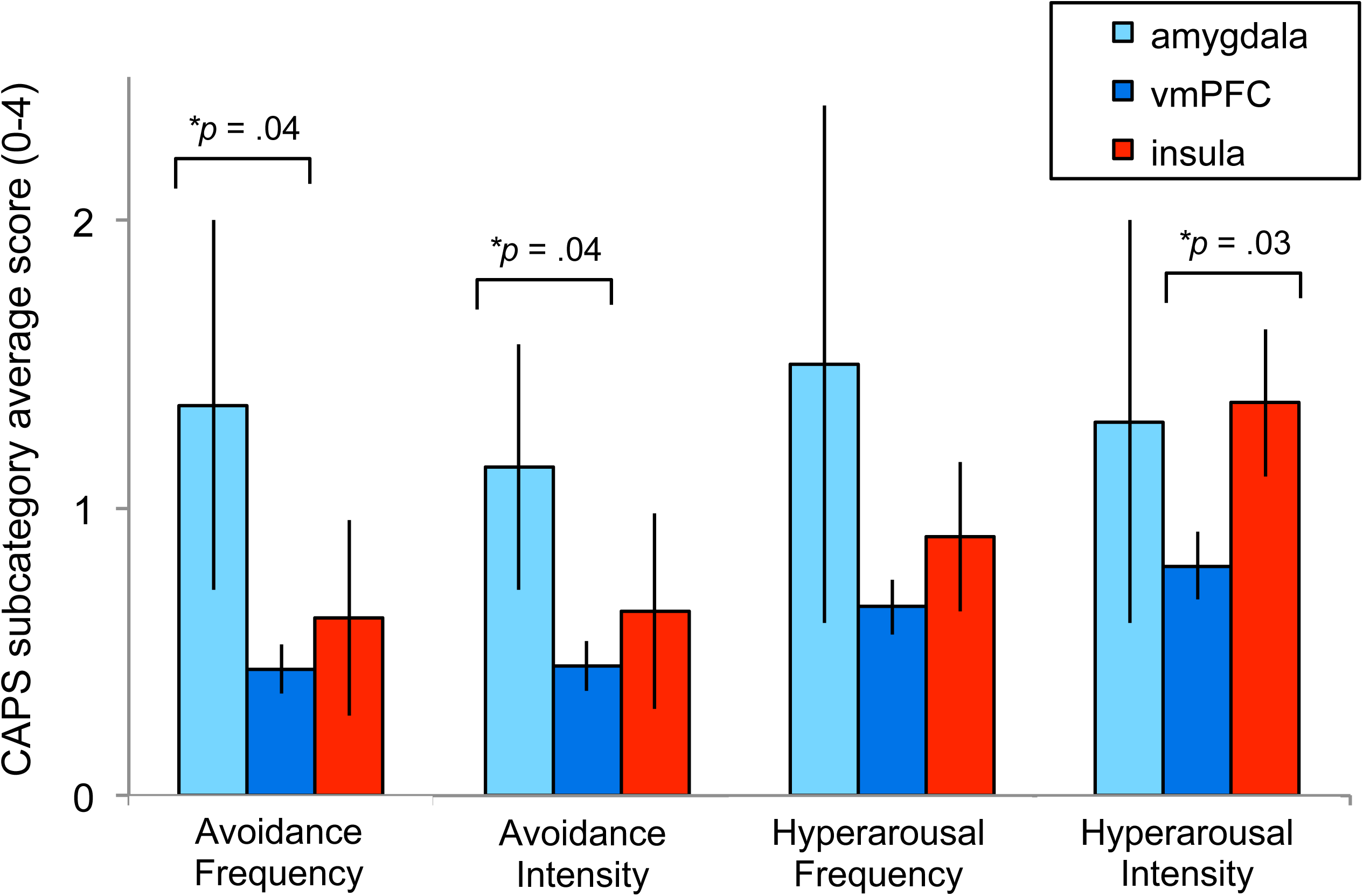
Avoidance and hyperarousal symptom scores for patients with lesions in amygdala, vmPFC, and insula. Patients with lesions predominantly in the amygdala showed significantly higher scores on avoidance frequency and avoidance intensity than those with lesions in vmPFC. Patients with lesions predominantly in the insula showed significantly higher hyperarousal intensity scores than patients with lesions in vmPFC. Note that neither re-experiencing scores nor scores from patients with hippocampus lesions are shown here, due to the lack of significant findings for these variables. Error bars signify SEM.

### Comparisons with Control Participants

Additional analyses were conducted to determine whether the effects noted above were specific to patients with lesions or whether they might be maintained in comparison to healthy subjects. Patients in lesion groups were thus compared to a matched control group of veterans (also drawn from the VHIS registry) with no history of brain injury (*n* = 54). For comparisons between patients with vmPFC lesions and healthy control participants, patients consistently showed decreased hyperarousal and avoidance: avoidance frequency (*U* = -449.5, *P* = 0.03); avoidance intensity (*U* = -455.5, *P* = 0.03); hyperarousal frequency (*U* = -467.0, *P* = 0.04); hyperarousal intensity (*U* = -432.5, *P* = 0.02). In contrast, group comparisons of re-experiencing symptoms showed no significant associations (all *P* > 0.77), nor did group comparisons between healthy control participants and all other lesion groups (amygdala, insula, hippocampus) on any symptom cluster (all *P* > 0.20). These results lend further support to the notion that vmPFC lesions are specifically and robustly associated with decreased avoidance and hyperarousal symptoms of PTSD.

Notably, analyses reported above did not yield results supporting a link between hippocampal lesions and re-experiencing symptoms, as hypothesized. Because the hippocampal lesion group had significantly lower lesion size variance than any other group (all *P* < .03), we conducted a post hoc analysis in which participants with any size hippocampal lesion (*N* = 25, Table 2) were compared to brain injured patients with no hippocampal lesions, thereby maximizing statistical variance without modifying selectivity criteria for the patient group. Due to increased statistical variance, non-significant Shapiro-Wilk tests supported use of parametric testing. Results revealed decreased re-experiencing symptoms among patients with hippocampal lesions in comparison to patients with no hippocampal lesions, for both symptom frequency (*t*_172_ = -2.09, *P* = .04) and intensity (*_t_*_172_ = -2.34, *P* = .02). In addition, patients with hippocampal lesions also reported fewer symptoms of avoidance, again for both frequency (*t*_172_ = -2.03, *P* = .04) and intensity (*t*_172_ = -2.09, *P* = .04) of symptoms. Importantly, results cannot be attributed to overall brain volume loss, which did not differ significantly between these two groups (*P* > .09). When patients with hippocampal lesions were further compared to the group of healthy participants (with no brain lesions) defined above, these results were maintained. Specifically, patients with hippocampal lesions showed decreased re-experiencing and avoidance symptoms on both measures of frequency and intensity (all *P* < .04). Given that these results were derived from patients who were not excluded for lesion volume, our findings suggest that links between the hippocampus and re-experiencing/avoidance symptoms may be sensitive even to relatively small changes in lesion volume.

### Effects of Potential Confounding Variables

Finally, to evaluate whether results might be influenced by confounding demographic or neuropsychological variables, we compared our sample of TBI participants to the matched control group of healthy participants described above (i.e., veterans with no history of brain injury). As shown in Table 1, TBI participants did not differ from healthy control participants in age or level of education. Regarding neuropsychological performance, TBI and healthy control participants showed comparable levels of pre-injury intelligence; however, those with brain injuries demonstrated lower post-injury intelligence (*t*_325_ = 3.97, *P* = 0.00009; see Table 1), as expected. Nevertheless, both groups’ scores were in the normal range for this measure. Additional measures of depression [Beck Depression Inventory-II; 51] and state/trait anxiety [State-Trait Anxiety Inventory; 52] did not differ between participants with TBI and those without. While patients in the control group showed slightly increased PTSD severity in comparison to patients with TBI, both in terms of symptom frequency and symptom severity [Clinician-Administered PTSD Scale; 11], these differences were not significant (all *P* > 0.19). Of note, our prior findings using this patient sample have demonstrated that TBI in key regions may act as a protective factor against maintaining PTSD diagnosis [19], indicating that trends in the direction of reduced PTSD among TBI patients should not be unexpected. Finally, In terms of combat experience, TBI participants were deployed for approximately 5 months less than healthy control participants (*t*256 = 5.45, *P* < 0.0001), due to brain injuries terminating their period of service. In addition, level of combat exposure was higher for TBI participants than for healthy control participants (*t*186 = 4.57, *P* < 0.0001).

Using the same set of demographic and neuropsychological variables, we also tested differences between the four lesion groups (amygdala, vmPFC, insula, hippocampus), as shown in Table 2. Paired comparisons were conducted between all combinations of lesion groups using Wilcoxon Mann Whitney tests, as described above. Comparisons yielded no significant results, indicating that the between-group findings reported above are not likely due to differences in the demographic and neuropsychological measures reported herein.

## Discussion

The purpose of the present study was to examine the effects of focal brain damage on individual PTSD symptom clusters, in order to determine whether key brain regions are causally involved in the heterogeneous expression of this disorder. Results from a unique sample of adults who experienced both traumatic brain injury and psychological trauma revealed that the amygdala, vmPFC, and hippocampus play a causal role in the pathogenesis of PTSD symptom clusters. Specifically, we report evidence for two distinct mechanisms that underlie symptom clusters of PTSD: (1) an amygdala-vmPFC network underlying hyperarousal and avoidance, and (2) a hippocampal network underlying re-experiencing and avoidance, as discussed below.

As hypothesized, amygdala damage was associated with hyperarousal symptoms, with decreased amygdala volume linked to increased symptoms of hyperarousal. Our findings replicate those reported in PTSD patients without TBI [33], suggesting that this result may be generalizable across both TBI and non-TBI populations. Our findings are also in alignment with research involving patients with non-traumatic focal brain damage to the amygdala (a rare genetic condition known as Urbach-Wiethe disease), who reliably exhibit hypervigilance across a variety of fear-inducing tasks [63]. The relationship between Urbach-Wiethe disease and the specific symptom of increased hypervigilance is of note, because patients with this disorder are well known to lack generalized fear responses. That is, patients with Urbach-Wiethe disease show hypervigilance but otherwise no symptoms of fear, even in situations that are objectively intensively fear-inducing. Critically, this phenomenon not only aligns with our present findings linking amygdala damage and increased hypervigilance but also with our previous research indicating that amygdala damage serves as a protective factor against PTSD in general [19], thus underscoring the complexity and specificity of the amygdala’s role in fear circuitry and PTSD. While the present study did not examine amygdala subregions, it is possible that our results may be supported, in part, by the burgeoning literature demonstrating that subnuclei of the amygdala relate to PTSD and fear-related behavior in distinct ways, as described above [35].

Numerous studies reporting on PTSD symptoms and regional activity have reported associations between increased hyperarousal symptoms and increased amygdala hyperreactivity [e.g., 35, 37, 64]. While the present study focused on changes in regional volume rather than changes in regional activity, recent rodent studies examining the relationship between focal injury and subsequent activity changes may offer further insight. These studies have demonstrated that structural damage may cause an initial period of regional hypoactivity, followed by subsequent longer term hyperactivity due to homeostatic activity regulation [65]. It is thus possible that volume loss in the amygdala may lead to an overcompensation of blood flow in this region, thereby resulting in amygdala hyperreactivity and increased hyperarousal symptoms.

Amygdala damage was also associated with avoidance symptoms, again with decreased amygdala volume correlating with increased symptoms. These results are in alignment with rodent studies that have demonstrated causal links between volume loss in the amygdala and increased fear avoidance behavior [35], with lesions of the central amygdala, specifically, leading to increased active avoidance (versus passive avoidance or freezing)[66]. To our knowledge, the present study may be the first to report the same causal link in humans, thus providing a bridge between rodent models of avoidance symptoms and their human counterpart. Findings from the present study also provide support for a recent hypothesis that redefines the role of the amygdala in PTSD as a decisional gateway for “behavioral engagement” with the feared stimulus [35] According to this hypothesis, disrupted amygdala circuitry specifically leads to behavioral avoidance of the feared stimulus. While this model has been argued to represent PTSD symptoms more accurately than traditional models based on Pavlovian fear learning [35], it lacks support in the human literature (partly due to the relatively small number of studies using methods that allow for testing of causal relationships between amygdala structure and function).

The present study also observed that vmPFC volume loss was associated with hyperarousal symptoms; critically, however, the direction of the association was reversed. That is, while decreased amygdala volume was associated with *increased* symptoms of hyperarousal, decreased vmPFC volume was associated with *decreased* symptoms. In addition, patients with vmPFC lesions reported decreased hyperarousal in comparison to patients with insular lesions and participants with no lesions. Although previous studies have reported contrasting findings regarding vmPFC activity and hyperarousal symptoms [37, 39], several studies have implicated this region in physiological arousal. Of particular relevance, vmPFC damage has been shown to blunt physiological arousal responses to fear-inducing stimuli [67–70]. Thus, if vmPFC plays a role in the expression of arousal, damage to this region may result in blunted responses of physiological arousal as well as decreased hyperarousal behavior. In this sense, vmPFC damage may act as a protective factor against hyperarousal symptoms as well as avoidance symptoms, given our additional results revealing reductions in this PTSD symptom cluster among participants in the vmPFC lesion group. Thus, taken together, findings from the present study support contrasting roles for amygdala and vmPFC, where damage to the amygdala acts as a risk factor for increased hyperarousal and avoidance, while vmPFC damage acts as a protective factor against the same symptoms.

In addition to findings regarding vmPFC and amygdala, our data revealed a unique role for the hippocampus in PTSD symptomatology. Our results indicated that patients with hippocampal lesions showed reduced re-experiencing and avoidance symptoms, in comparison to both TBI patients with no hippocampal lesions and healthy participants with no TBI. Prior research on the role of the hippocampus in PTSD symptom clusters is scant, with one study reporting an association between dysregulated hippocampal resting-state connectivity and increased re-experiencing symptoms [40]. However, research on the role of the hippocampus in PTSD as a unitary construct is profuse [13, 16, 17], with numerous studies reporting low hippocampal volume in PTSD patients [13, 26, 27], as well as abnormal activity during associative memory and fear-related tasks [29–32]. While hippocampal involvement in PTSD is well established in this area of literature, the precise nature of the hippocampus’ role in this disorder remains undefined. It has been suggested that this role likely involves contextual memory dysfunction [e.g., 13], considering that contextual memory is a key function of the hippocampus and that PTSD involves deficits in applying contextual cues to signal safety. While this theory remains largely speculative, our finding relating the hippocampus to re-experiencing symptoms may provide a new and critical source of support, given that the re-experiencing symptom cluster includes items descriptive of contextual memory deficits (e.g., “acting or feeling as if event were recurring”). Thus, our findings may not only contribute key evidence that the role of the hippocampus in PTSD involves contextual memory deficits, but furthermore, our methodology suggests that the role of the hippocampus in the pathogenesis of these deficits is causal.

Despite the unique opportunity to sample a large set of patients with focal brain lesions, we acknowledge specific study limitations. First, our sample was homogenous in terms of gender, age, and trauma source. Therefore, it may be difficult to interpret how results might generalize to a more diverse population, particularly given that gender [71, 72] and trauma source [73] have been found to play a role in the neural mechanisms underlying PTSD. Furthermore, our measure of lifetime PTSD (as opposed to current PTSD) includes retrospective self report (i.e., CAPS), which could be concerning in population of TBI patients, potentially weakening the validity of their responses [74, 75]. Finally, although all participants in the present study had been exposed to trauma and had experienced PTSD symptomatology (participants with no history of symptoms were excluded), symptom severity and frequency were relatively mild, and the majority of our sample did not meet criteria for diagnosis. It is possible that a participant sample more representative of full PTSD diagnosis would present with a distinct pattern of focal damage in relation to symptom clusters; however, recent research suggests that differing diagnostic categories of PTSD (based on symptom severity) are not associated with structural differences between brain regions [76]. Rather, a more severe PTSD diagnosis was associated with structural differences *within* affected regions (with more severe diagnostic classification associated with decreased volume in entorhinal cortex and rostral ACC), potentially suggesting that the strength of our findings may be amplified in a sample with a higher number of PTSD cases.

These limitations notwithstanding, the present study is the first to reveal effects of a brain damage on PTSD symptom clusters, thus providing evidence for causal roles of amygdala and vmPFC in hyperarousal and avoidance symptoms, as well as the hippocampus in symptoms of re-experiencing and avoidance. To further probe the roles of these brain regions in PTSD symptomatology, future studies might examine whether methods of targeting regional brain activity, such as non-invasive brain stimulation, might be effective in modifying behaviors associated with particular PTSD symptom clusters. With a better understanding of the brain regions involved in distinct symptom clusters, we will not only develop a more complete picture of the neural mechanisms underlying PTSD, but also aid in the development of individualized therapeutic strategies that could be used to target distinct symptom profiles [12].

## Acknowledgments and Disclosures

The authors would first like to thank the Vietnam veterans who donated their time and energy to participating in this study. We also thank the National Institute of Neurological Disorders and Stroke and the National Naval Medical Center for providing facilities and supporting this research. Finally, we thank G. J. Solomon, V. Raymont, S. Bonifant, B. Cheon, C. Ngo, A. Greathouse, K. Reding, and G. Tasick for the testing and evaluation of participants. Support was provided by the Therapeutic Cognitive Neuroscience Fund to B. G., the Agency for Healthcare Research and Quality to K. K. H. [K12 HS023011], the Dixon Translational Research Grants Initiative to K. K. H., and the Brain and Behavior Research Foundation’s NARSAD Young Investigator Award to K. K. H.

All authors report no biomedical financial interests or potential conflicts of interest.

